# Sex-Dependent Effects of Creatine Hydrochloride on Neuroinflammation, Neuronal Survival, and Inhibitory Synaptic Remodeling Following Traumatic Brain Injury in Mice

**DOI:** 10.64898/2026.10.21.760461

**Authors:** Johnny W. Zigmond, Benjamin G. Lamberty, Gretchen J. Simon, Jonathan M. Wood, Natalie M. Elliott, Don W. Miller, Kelly L. Stauch

**Affiliations:** University of Nebraska Medical Center, College of Medicine, Department of Neurological Sciences, Omaha, NE, USA; University of Manitoba, Department of Pharmacology and Therapeutics, Winnipeg, Canada

## Abstract

Despite traumatic brain injury (TBI) being a leading cause of death and disability in the United States, there is currently no FDA-approved medication that can prevent, heal, or reverse brain damage caused by TBI. TBI is caused by primary and secondary brain damage resulting in loss of neurons, astrocytic gliosis, and microglial activation which culminate in complex neurological disorders. The goal of the current study was to investigate the neuroprotective potential for Creatine Hydrochloride (C-HCl) supplementation following TBI in wild-type mice. Four C-HCl supplementation paradigms were examined including both 7 day pre-injury and 14-day post-injury treatments consisting of the following: 1) water/water, 2) water/C-HCl (100 mg/kg), 3) C-HCl (100 mg/kg)/C-HCl (100 mg/kg), 4) C-HCl (100 mg/kg)/C-HCl (450 mg/kg). Following controlled cortical impact (CCI) injury at a parietal location, measures of tissue and cellular (neuronal and glial) damage were assessed at 14 days post-injury (dpi). At 14 dpi, there were no significant effects on gross anatomical cavitation, tissue loss, third ventricle expansion, or systemic fluid biomarkers with any treatment group, though female mice exhibited larger baseline lesion boundaries overall compared to male mice. However, microstructural evaluation revealed a profound and sex-dependent effect of C-HCl supplementation in the hippocampus with female mice showing reduced neuroinflammation and male mice displaying reduced neurodegeneration (Fluoro-Jade C staining of neurons) across the CA1 and dentate gyrus (DG) compared to placebo control. Furthermore, mice receiving C-HCl pre-injury followed by high-dose C-HCl post-injury demonstrated distinct remodeling of inhibitory circuits to restore network balance with males significantly modifying presynaptic machinery via vesicular GABA transport (VGAT puncta) in the DG, whereas females reinforced postsynaptic architectural framing through scaffolding elements (gephyrin puncta) in both the CA1 and DG. These results suggest C-HCl supplementation holds promise as a neuroprotective therapeutic strategy for TBI. Crucially, the molecular mechanisms utilized to achieve this brain cell homeostasis are radically distinct between the sexes, females benefit primarily via upstream glial stabilization, while males derive protection via direct downstream neuronal survival. These outcomes highlight a critical requirement for sex-stratified profiling in translational pipelines for neurometabolic interventions for TBI. Ultimately, these data indicate that C-HCl successfully counteracts chronic secondary injury cascades, mitigating neurodegeneration, dampening neuroinflammation, and remodeling inhibitory circuitry to prevent post-traumatic network hyperexcitability.

## Introduction

Creatine (N-[aminoiminomethyl]-N-methyl glycine) is an amino acid-like compound that plays a central role in cellular energy metabolism ^1^. Creatine and the creatine kinase/phosphocreatine system play an important role as reserved sources of energy in tissues with high and fluctuating energy requirements. The role of creatine in energy metabolism involves the transfer of N-phosphoryl groups from phosphorylcreatine to adenosine diphosphate (ADP) to regenerate adenosine triphosphate (ATP) through a reversible reaction catalyzed by phosphorylcreatine kinase. Creatine from either dietary or endogenous sources are distributed via peripheral circulation to tissues with approximately 95% of the total creatine pool stored in skeletal muscle, while the remaining 5% is distributed amongst other metabolically demanding organs, such as the brain, heart, and kidney. Historically, research on creatine has focused primarily on skeletal muscle; however, over the past decade there has been an increased appreciation for the potential value of creatine supplementation for supporting brain health and cognitive function ^1,2^. There is growing evidence that creatine may be of value in the treatment of a number of neurological and psychiatric disorders.

Traumatic brain injury (TBI), a leading cause of disability and death in the United States, induces both focal damage as well as more diffuse disruption in neural function ^3^. The primary injury occurs due to the direct mechanical forces acting upon the brain tissue, while secondary injury evolved over time and results from the cellular and molecular processes initiated by the initial trauma, including cerebral edema, glial and neuronal dysfunction, inflammatory signaling, and metabolic alterations. Based on results of both animal and human studies ^2^, creatine represents a promising nutritional supplement for increasing resilience to and treating TBI.

Controlled cortical impact (CCI) is a commonly used and highly regarded model of brain trauma that uses a pneumatically or electromagnetically controlled piston to induce a reproducible and well-controlled injury ^4^. Animal models of TBI are of immeasurable value to better understand outcomes of brain trauma and in testing potential therapeutic agents. While creatine treatment holds promising translational value for improving brain health, studies on creatine supplementation in the context of TBI remain limited. This study aims to fill this gap by evaluating the neuroprotective effects of Creatine Hydrochloride (C-HCl) supplementation when administered after or prior to and after TBI. Mice were divided into four treatment groups: 1) water for 7 days prior to and 14 days after CCI, 2) water for 7 days prior to then 100 mg/kg C-HCl for 14 days after CCI, 3) 100 mg/kg C-HCl for 7 days prior to then 100 mg/kg C-HCl for 14 days after CCI, 4) 100 mg/kg C-HCl for 7 days prior to then 450 mg/kg C-HCl for 14 days after CCI; with post-injury treatments starting 30 min after surgery completion. The impacts of C-HCl treatment on tissue damage, blood biomarkers, glial activation, neuron degeneration, and synapse loss were investigated. While neither male nor female mice showed changes in TBI-induced tissue damage when supplemented with C-HCl/ C-HCl (450), the number of degenerating neurons found in the hippocampus were significantly lower in C-HCl/C-HCl (450) treated male mice. Further, the numbers of synaptic puncta were significantly higher in the C-HCl/C-HCl (450) treated male and female mouse hippocampus. Reduced astrocytic gliosis and microglial activation were observed in the hippocampus of female mice treated with C-HCl/C-HCl (450) and while not significant in male mice similar trends were observed. These findings indicate that while C-HCl therapy fails to alter macroscopic tissue pathology, it drives targeted, sub-cellular neuroprotection and restores inhibitory networks within the injured hippocampus in a sex-dependent manner with females demonstrating reduced glial activation and males exhibiting neuronal survival.

## Methods

### Animals

Male and female C57BL/6NCrl (from Charles River Laboratories) wild-type (WT) mice were randomly assigned to the four treatment groups. TBI surgeries were performed when the mice were 2 months of age. Mice were sacrificed at 14 days post-injury (dpi). Following surgery, mice were single-housed in a temperature-controlled environment with 12-hour light/dark cycle with free access to food and water at the Department of Comparative Medicine at the University of Nebraska Medical Center (UNMC). Experiments were approved and conducted according to the UNMC Institutional Animal Care and Use Committee (IACUC).

### Creatine Hydrochloride (C-HCl) treatment

C-HCl was obtained from Vireo Systems. The C-HCl was prepared fresh daily in sterile water and administered to the mice daily via oral gavage. The four treatment groups included: 1) water for 7 days prior to TBI and 14 days after TBI starting 30 min after surgery completion (water/water), 2) water for 7 days prior to TBI then 100 mg/kg C-HCl for 14 days after TBI starting 30 min after surgery completion (water/C-HCl), 3) 100 mg/kg C-HCl for 7 days prior to TBI then 100 mg/kg C-HCl for 14 days after TBI starting 30 min after surgery completion (C-HCl/C-HCl), 4) 100 mg/kg C-HCl for 7 days prior to TBI then 450 mg/kg C-HCl for 14 days after TBI starting 30 min after surgery completion (C-HCl/C-HCl (450)). Using the human equivalent dose calculations based on body surface area ^5^, the mouse doses of 100 and 450 mg/kg would be equal to 8.13 and 36.59 mg/kg (animal dose in mg/kg divided by 12.3 equals human equivalent dose), which when using the reference body weight for a human (60 kg) equals 487.8 and 2,195.4 mg (or 0.487 and 2.1954 g).

### Controlled cortical impact (CCI)

Two-month-old mice were anesthetized using 2.5% isoflurane through inhalation in an anesthesia induction box. Their heads were shaved and they were placed in a stereotaxic frame (RMD) where they were maintained on isoflurane delivered by a nose cone. A midline incision was made to expose the skull, and a 3.5 mm craniotomy was made using a trephine and Dremel centered at -2.5 mm bregma and 2.75 mm lateral to the midline over the left parietal cortex. Mice were subjected to CCI injury at a 0.5 mm depth, 3.5 m/s speed, and 200 ms dwell time. The 68099 II Precision Impactor Device (RWD) uses a computer controlled pneumatically driven piston to rapidly impact the brain. After injury, the incision was closed using wound clips and the mice were administered 1 mg/kg of buprenorphine ER (Epicur Pharma) by subcutaneous injection for pain and allowed to recover on a heating pad. Mice were then given appropriate C-HCl treatment or vehicle control (water) by oral gavage 30 min post-surgery and every subsequent 24 h for 14 days.

### Injury area fraction

Once the brains were gently dissected, prior to histology tissue processing, photographs of the whole dorsal brain were taken to analyze the injury area fraction. For analysis, ImageJ was used to outline an ROI around the injury, and its area was measured and normalized to the area of the contralateral hemisphere to obtain an injury area fraction.

### Brain processing and sectioning

Brains were gently removed and fixed in 4% paraformaldehyde in 1x PBS for 24 h at 4°C. After fixation, brains were rinsed with 1x PBS three times for 10 min prior to cryoprotection using 30% sucrose in 1x PBS for 48 h at 4°C. After cryoprotection, the brains were rinsed with 1x PBS three times for 10 min prior to embedding in 3% agar in 1x PBS for tissue sectioning using a vibrating microtome (Leica). Coronal sections were placed on Super Frost Plus microscope slides and stored at -20°C until staining.

### Hematoxylin and eosin slide scanner

50 μm thick coronal sections were sent to the UNMC Tissue Core Facility for processing and staining with hematoxylin and eosin (H&E) followed by imaging under a slide scanner (Leica Aperio CS2 brightfield) at 40x magnification power and analyzed using QuPath software. For analysis, an ROI annotation was drawn around the contralateral hemisphere, ipsilateral hemisphere, ipsilateral hippocampus, and third ventricle. The are for each structure was measured and normalized to the area of the contralateral hemisphere to obtain an area fraction.

### Immunohistochemistry

Coronal sections were stained as previously described by our group ^6^. In brief, 50 μm thick coronal sections were stained with the following primary antibodies: GFAP (1:1000, Cat ab68428, Abcam), Iba1 (1:1000, Cat #234009, Synaptic Systems), PSD95 (1:500, Cat #124008, Synaptic Systems), VGLUT1 (1:500, Cat #135316, Synaptic Systems), Gephyrin (1:500, Cat #147318, Synaptic Systems), and VGAT (1:500, Cat #131008, Synaptic Systems). For secondary antibody conjugation, the following antibodies were used: Alexa Goat anti-Rabbit 568 (1:1000, Cat #A11011, Invitrogen), Alexa Goat anti-Chicken 647 (1:1000, Cat #A21449, Invitrogen), Alexa Goat anti-Guinea Pig 647 (1:1000, Cat #A21450, Invitrogen), and DAPI for nuclear counterstain.

### FluoroJade staining

Coronal sections were stained with Fluoro-Jade C (Cat #NC0859611, Fisher Scientific) according to the manufacturer’s instructions. Briefly, slides were prepared with sodium hydroxide, 70% ethanol, and potassium permanganate solutions. Fluoro Jade C stain was coupled with DAPI Nissl counterstain and incubated in a dark environment at room temperature for 10 minutes. After drying on a slide warmer (50-60℃) for 5 minutes, slides were submerged in xylenes, and DPX mounting media was used to place the coverslip.

### Fluorescent confocal imaging and image quantification

Stained slides were imaged at 40x-oil immersion using a Nikon A1R upright confocal microscope. Images were obtained at 1024 x 1024-pixel size with 561 and 640 nm lasers. Stained slides were imaged at 0.5 μm steps with 21-29 steps per z-stack. Analysis was performed in ImageJ for quantification of percent (% area), mean fluorescence intensity (MFI), and puncta counts from the corresponding 40x images in three different ipsilateral brain regions: Cornu Ammonis 1 (CA1) and Dentate Gyrus (DG) regions of the hippocampus and the parietal cortex.

### Statistical analysis

Statistical analyses were performed using GraphPad Prism 11 software and outliers were identified using Grubbs’ test with α = 0.05. One-way ANOVA was used to compare the effect of treatment group, which if significant was followed by Tukey’s multiple comparisons test. Two-way ANOVA was used to compare the effect of sex and treatment, which if significant was followed by Sidak’s multiple comparisons test. In accordance with rigor and reproducibility, the investigators were blinded during experiments and analyses, and animals were randomly assigned to each cohort.

## Results

### Injury area

First, we assessed if C-HCl treatment had an impact on the TBI-induced lesion size and loss of hippocampal tissue of male and female mice at 14 dpi. Photographs of the dorsal brain were taken to measure the area fraction of the injury normalized to the contralateral hemisphere. C-HCl treatment did not significantly affect injury area size in mice of either sex at 14 dpi (**Figure 1**). The TBI-induced injury area was significantly greater in female compared to male mice regardless of treatment group (**Supplemental Figure S1**). Coronal brain sections were stained with hematoxylin and eosin (H&E) and imaged using a whole-slide scanner. The ipsilateral hemisphere area, ipsilateral hippocampal area, and third ventricle area were measured and normalized to the area of the contralateral hemisphere to uncover any tissue loss and/or ventricle enlargement. No significant changes were uncovered in any of these measures at 14 dpi between treatment groups (**Figure 2**).

**Figure 1.**
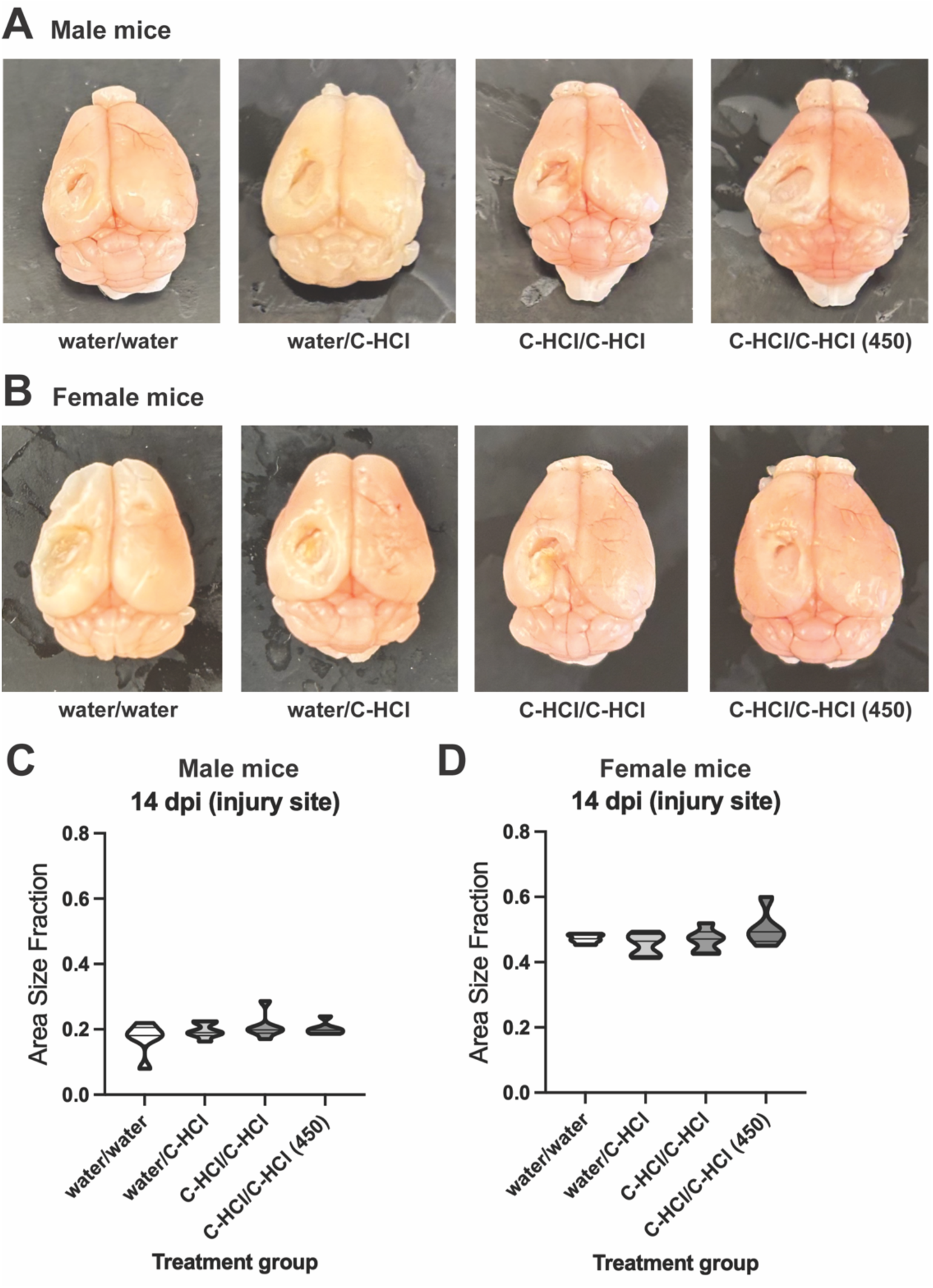
TBI-induced tissue damage is unaltered at 14 dpi in mice treated with C-HCl. Representative photographs of the dorsal brain from (A) male and (B) female mice treated with water/water, water/C-HCl, C-HCl/C-HCl, and C-HCl/C-HCl (450) for 7 days before/14 days after TBI. C-HCl concentration was 100 mg/kg unless noted as 450 mg/kg. The area fraction of the injury site (area of the injury site/area of the contralateral brain hemisphere) for (C) male and (D) female mice at 14 dpi. Data represented as a volcano plot of *n* = 6 per treatment group. No significant differences detected between treatment groups (one-way ANOVA).

**Figure 2.**
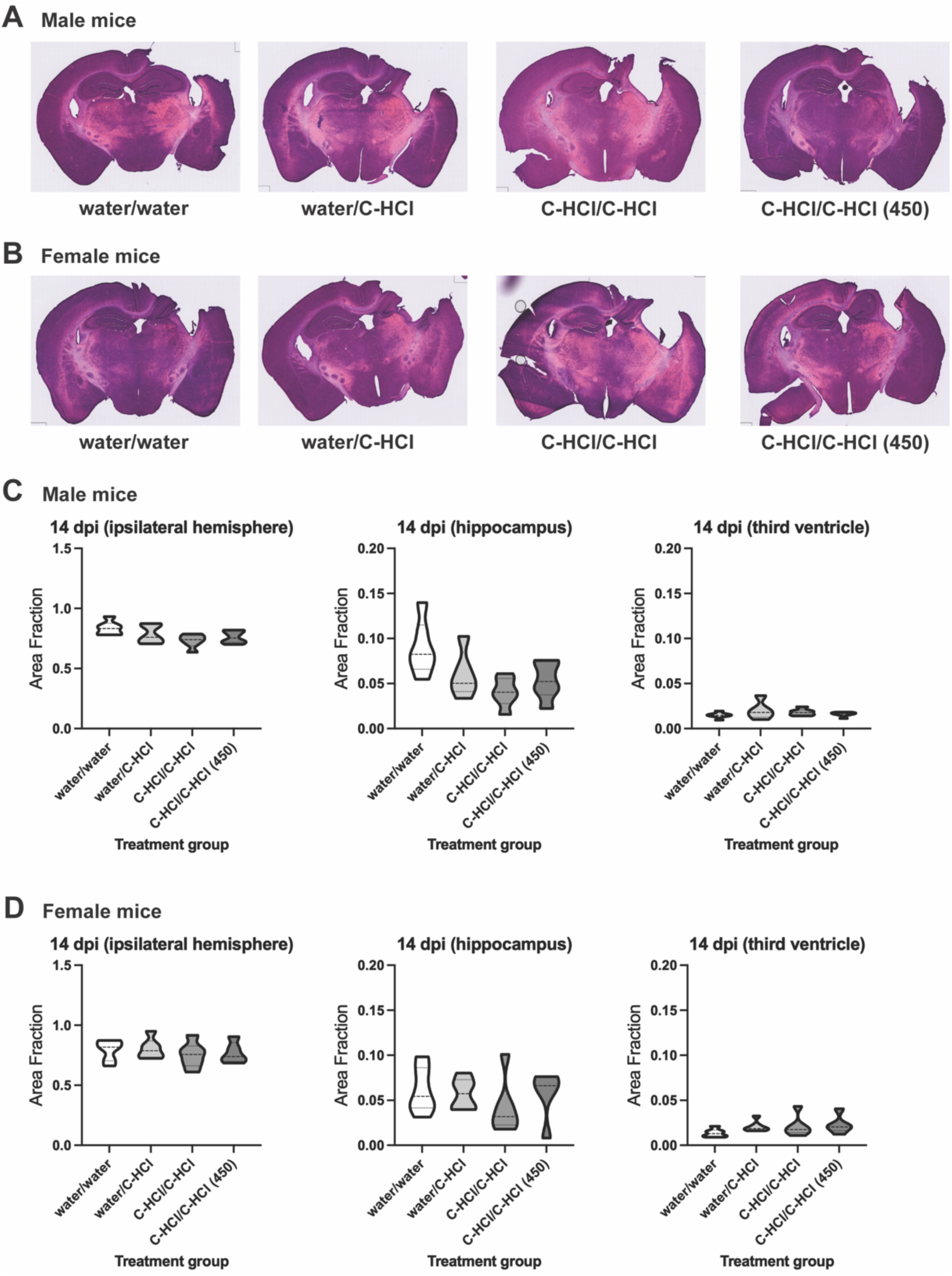
TBI-induced tissue degeneration is unaltered at 14 dpi in mice treated with C-HCl. H&E staining of coronal brain sections from (A) male and (B) female mice treated with water/water, water/C-HCl, C-HCl/C-HCl, and C-HCl/C-HCl (450) for 7 days before/14 days after TBI. C-HCl concentration was 100 mg/kg unless noted as 450 mg/kg. The area fraction of the ipsilateral hemisphere, ipsilateral hippocampus, and third ventricle for (C) male and (D) female mice at 14 dpi. Data represented as a volcano plot of *n* = 5-6 per treatment group. No significant differences detected between treatment groups (one-way ANOVA).

### Blood biomarkers

Neurofilament light chain (NfL) and total tau (t-Tau) are proteins released into the blood and cerebrospinal fluid (CSF) when neurons are damaged. Glial fibrillary acidic protein (GFAP) is a protein whose detection in the blood and CSF indicates astrocyte damage and blood-brain barrier disruption. At 14 dpi blood was collected prior to sacrifice via cardiac puncture, processed to obtain plasma, and analyzed using ELISAs to measure the levels of NfL, t-Tau, and GFAP. No significant changes were uncovered in any of these measures at 14 dpi between treatment groups (**Figure 3**).

**Figure 3.**
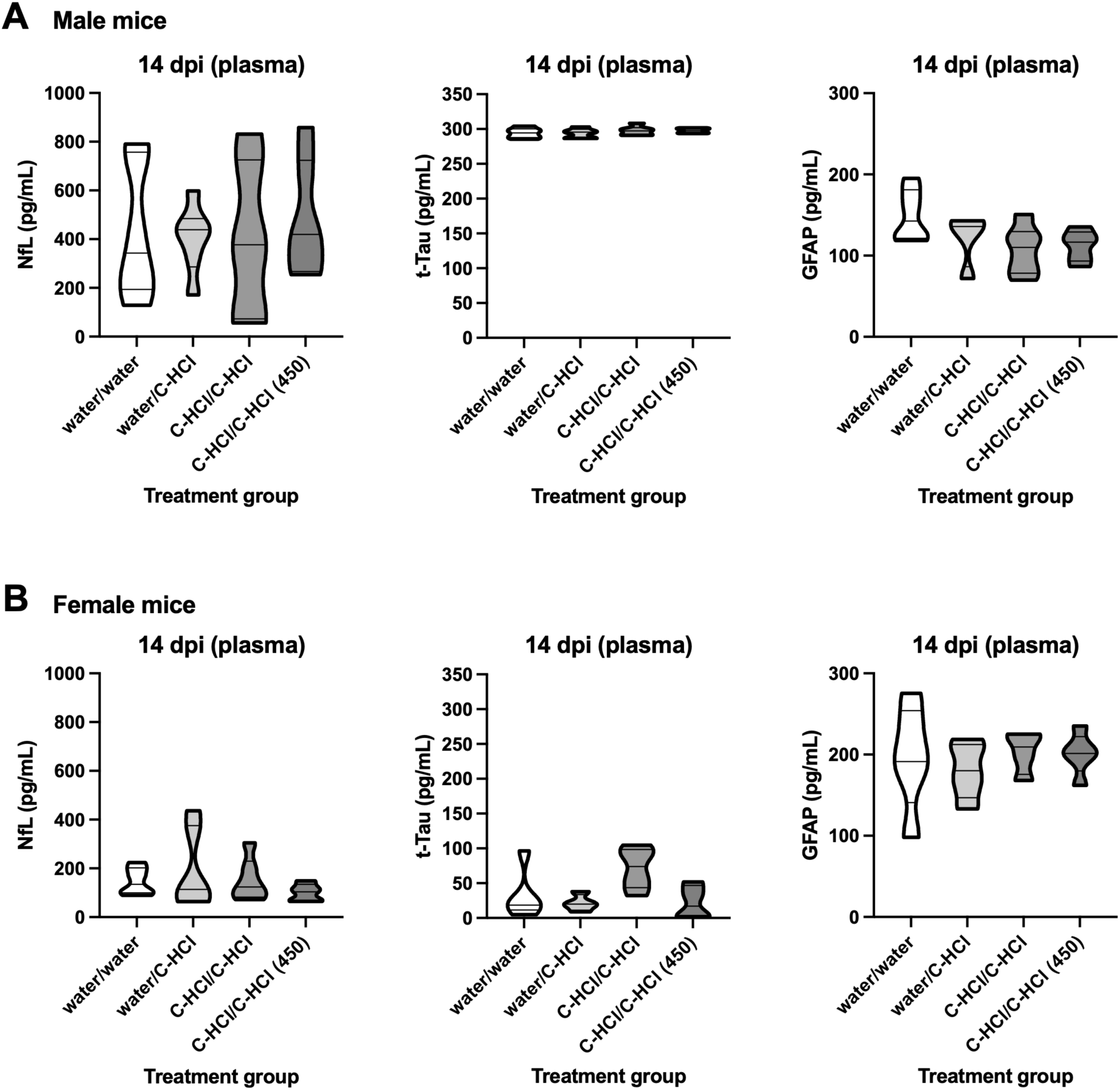
Plasma levels of NfL, t-Tau, and GFAP are unaltered at 14 dpi in mice treated with C-HCl. ELISA assay measurements of NfL, t-Tau, and GFAP in plasma from (A) male and (B) female mice treated with water/water, water/C-HCl, C-HCl/C-HCl, and C-HCl/C-HCl (450) for 7 days before/14 days after TBI. C-HCl concentration was 100 mg/kg unless noted as 450 mg/kg. Data represented as a volcano plot of *n* = 4-6 per treatment group. No significant differences detected between treatment groups (one-way ANOVA).

### Glial cells

We compared the GFAP and ionized calcium-binding adaptor molecule 1 (Iba1) staining between the treatment groups to evaluate astrocyte (GFAP) and microglial (Iba1) activation at 14 dpi. We examined the ipsilateral hippocampus (Cornu Ammonis 1 (CA1) and Dentate Gyrus (DG) regions) and parietal cortex. No significant changes in either microglia density (% area of fluorescent staining) or activation (mean fluorescent intensity (MFI) of staining) were observed across treatment groups in male mice at 14 dpi (**Figure 4**). Similarly, C-HCl treatment did not significantly affect either astrocyte density or activation in male mice at 14 dpi (**Figure 5**). C-HCl treatment resulted in trends towards lower microglial density and activation which reached significance for decreased microglial activation in the C-HCl/C-HCl (450) versus water/water treated female mice at 14 dpi in the CA1 region of the hippocampus (**Figure 6**). Similar effects were observed for astrocytes in the hippocampal CA1 region of female mice at 14 dpi with significantly reduced astrocyte density and activation in the C-HCl/C-HCl (450) versus water/C-HCl treatment groups (**Figure 7**). The female mice in the C-HCl/C-HCl compared to water/C-HCl treatment group exhibited significantly lower astrocyte activation at 14 dpi in the CA1 region of the hippocampus (**Figure 7**).

**Figure 4.**
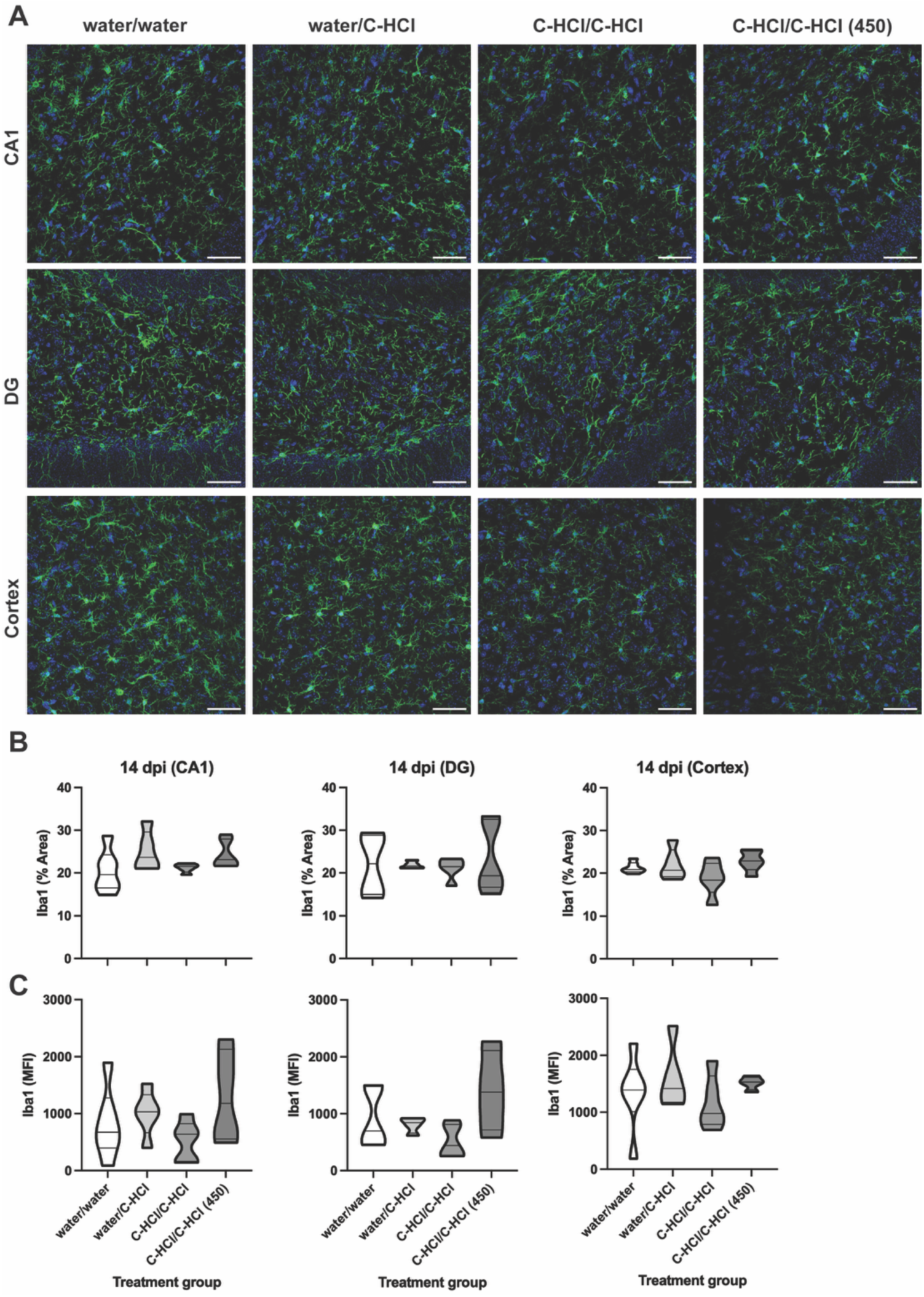
TBI-induced microglial activation is unaltered at 14 dpi in male mice treated with C-HCl. (A) Representative immunofluorescent images of Iba1 (green) and DAPI (blue) staining of coronal brain sections from male mice treated with water/water, water/C-HCl, C-HCl/C-HCl, and C-HCl/C-HCl (450) for 7 days before/14 days after TBI. C-HCl concentration was 100 mg/kg unless noted as 450 mg/kg. Objective/scale bar: 40x/50 μm. Quantification of the (B) percent (%) area and (C) mean intensity of the fluorescent staining in the CA1 hippocampus (left panel), DG hippocampus (middle panel), and parietal cortex (right panel). Data represented as a volcano plot of *n* = 4-6 per treatment group. No significant differences detected between treatment groups (one-way ANOVA).

**Figure 5.**
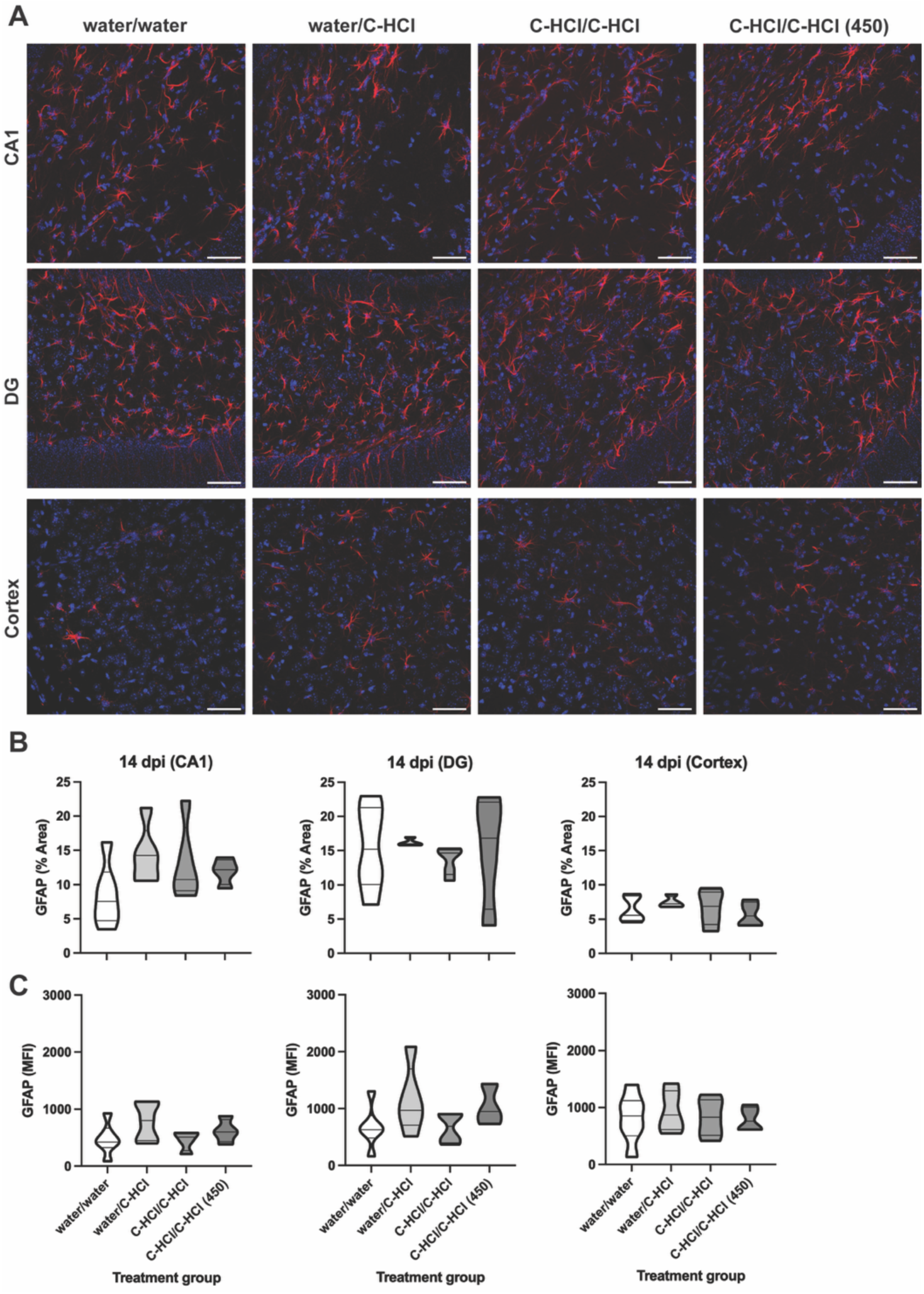
TBI-induced astrocyte activation is unaltered at 14 dpi in male mice treated with C-HCl. (A) Representative immunofluorescent images of GFAP (red) and DAPI (blue) staining of coronal brain sections from male mice treated with water/water, water/C-HCl, C-HCl/C-HCl, and C-HCl/C-HCl (450) for 7 days before/14 days after TBI. C-HCl concentration was 100 mg/kg unless noted as 450 mg/kg. Objective/scale bar: 40x/50 μm. Quantification of the (B) percent (%) area and (C) mean intensity of the fluorescent staining in the CA1 hippocampus (left panel), DG hippocampus (middle panel), and parietal cortex (right panel). Data represented as a volcano plot of *n* = 4-6 per treatment group. No significant differences detected between treatment groups (one-way ANOVA).

**Figure 6.**
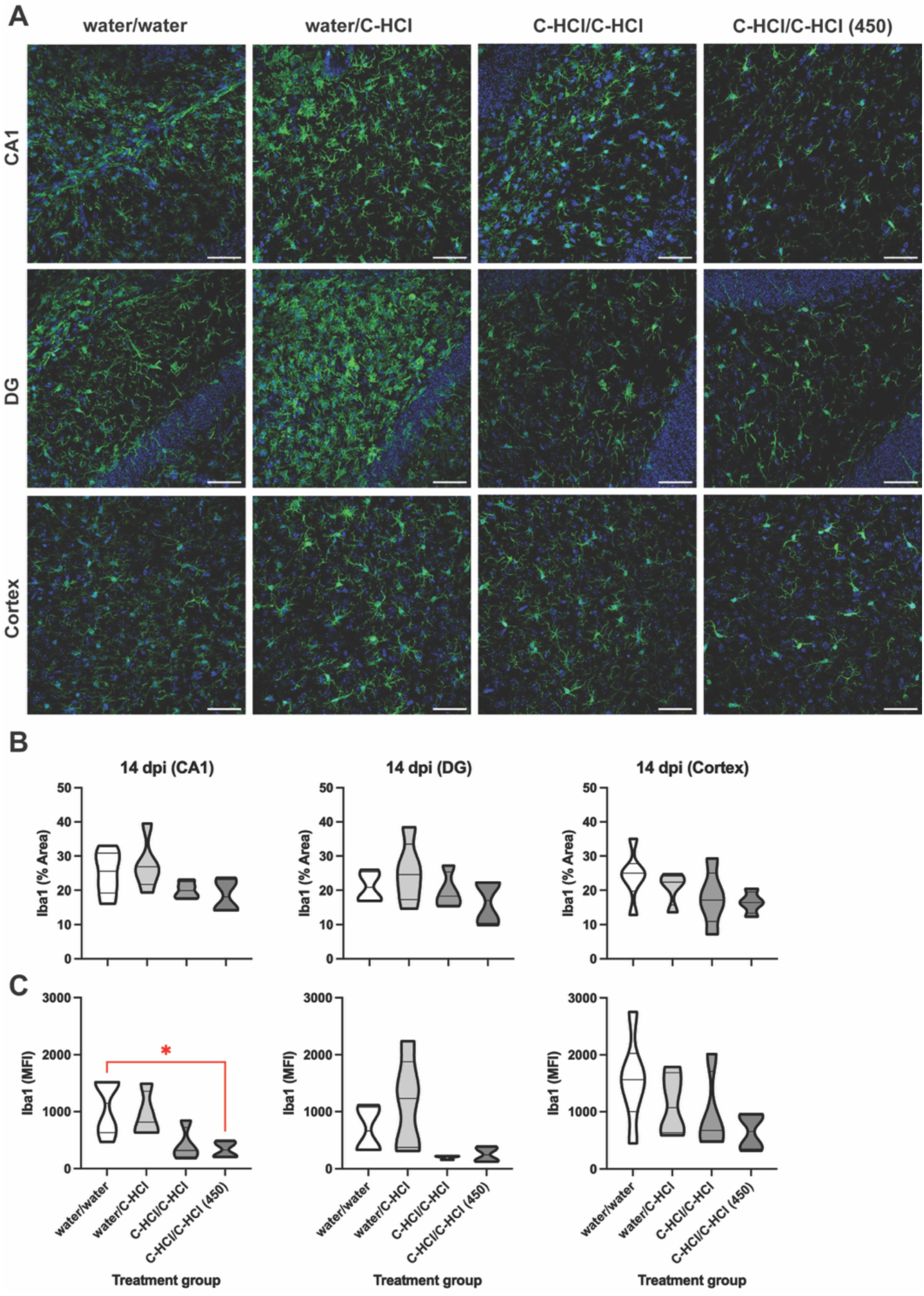
C-HCl treatment reduces microglial activation at 14 dpi in female mice. (A) Representative immunofluorescent images of Iba1 (green) and DAPI (blue) staining of coronal brain sections from female mice treated with water/water, water/C-HCl, C-HCl/C-HCl, and C-HCl/C-HCl (450) for 7 days before/14 days after TBI. C-HCl concentration was 100 mg/kg unless noted as 450 mg/kg. Objective/scale bar: 40x/50 μm. Quantification of the (B) percent (%) area and (C) mean intensity of the fluorescent staining in the CA1 hippocampus (left panel), DG hippocampus (middle panel), and parietal cortex (right panel). Data represented as a volcano plot of *n* = 4-6 per treatment group. Statistical significance was determined using a one-way ANOVA followed by Tukey’s multiple comparisons test. \**p* < 0.05.

**Figure 7.**
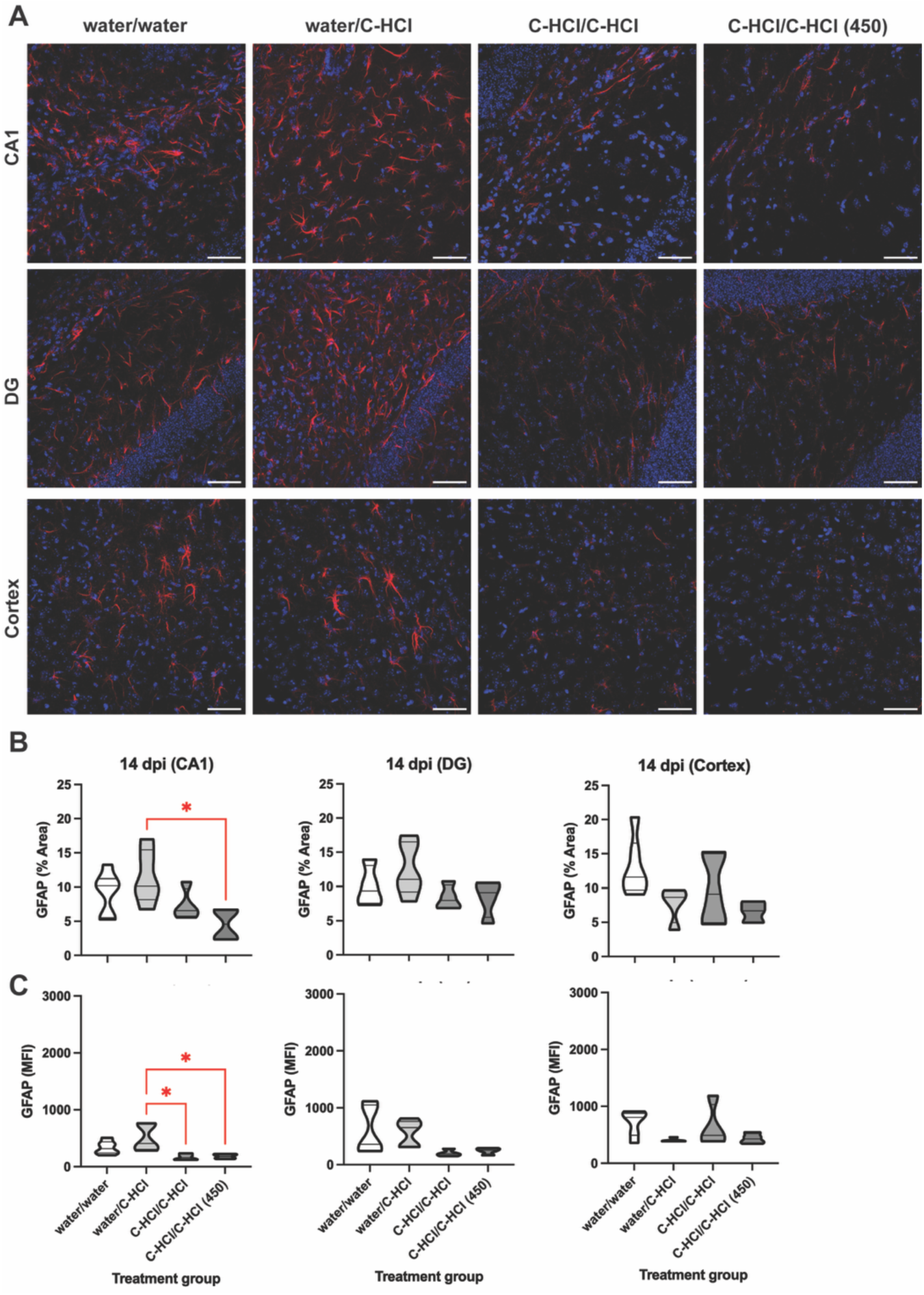
C-HCl treatment lowers astrocyte activation at 14 dpi in female mice. (A) Representative immunofluorescent images of GFAP (red) and DAPI (blue) staining of coronal brain sections from female mice treated with water/water, water/C-HCl, C-HCl/C-HCl, and C-HCl/C-HCl (450) for 7 days before/14 days after TBI. C-HCl concentration was 100 mg/kg unless noted as 450 mg/kg. Objective/scale bar: 40x/50 μm. Quantification of the (B) percent (%) area and (C) mean intensity of the fluorescent staining in the CA1 hippocampus (left panel), DG hippocampus (middle panel), and parietal cortex (right panel). Data represented as a volcano plot of *n* = 4-6 per treatment group. Statistical significance was determined using a one-way ANOVA followed by Tukey’s multiple comparisons test. \**p* < 0.05.

### Neuron loss

Fluoro-Jade C staining was used to label degenerating neurons at 14 dpi. We examined the ipsilateral hippocampal CA1 and DG regions and the parietal cortex. Overall, C-HCl treatment resulted in trends towards lower numbers of degenerating neurons with significant decreases in neuron degeneration observed in the C-HCl (450) versus water/water treated male mice at 14 dpi in the CA1 and DG regions of the hippocampus (**Figure 8**). Significantly reduced numbers of degenerating neurons were observed in the hippocampal DG region at 14 dpi in male mice treated with water/C-HCl versus water/water (**Figure 8**). While the absolute number of Fluoro-Jade C positive neurons were less in female mice following TBI compared to males, no significant changes in the presence of degenerating neurons (mean fluorescent intensity (MFI) of staining) were observed across treatment groups in female mice at 14 dpi (**Figure 9**).

**Figure 8.**
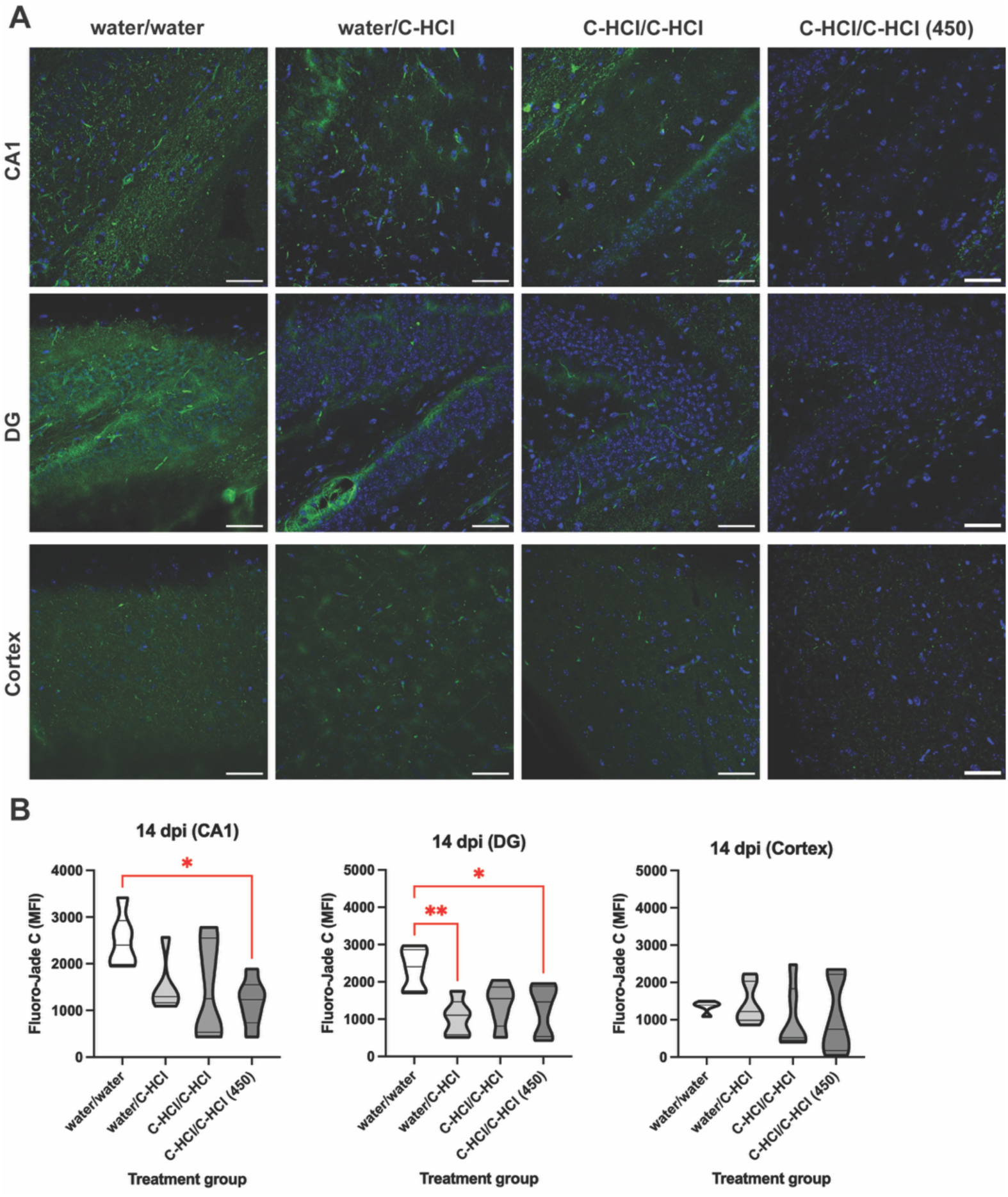
C-HCl treatment decreased neuron degeneration at 14 dpi in male mice. (A) Representative immunofluorescent images showing Fluoro-Jade C staining marking degenerating neurons in coronal brain sections from male mice treated water/water, water/C-HCl, C-HCl/C-HCl, and C-HCl/C-HCl (450) for 7 days before/14 days after TBI. C-HCl concentration was 100 mg/kg unless noted as 450 mg/kg. Objective/scale bar: 40x/50 μm. (B) Quantification of the mean intensity of the fluorescent staining in the CA1 hippocampus (left panel), DG hippocampus (middle panel), and parietal cortex (right panel). Data represented as a volcano plot of *n* = 5-6 per treatment group. Statistical significance was determined using a one-way ANOVA followed by Tukey’s multiple comparisons test. \**p* < 0.05, \*\**p* < 0.01.

**Figure 9.**
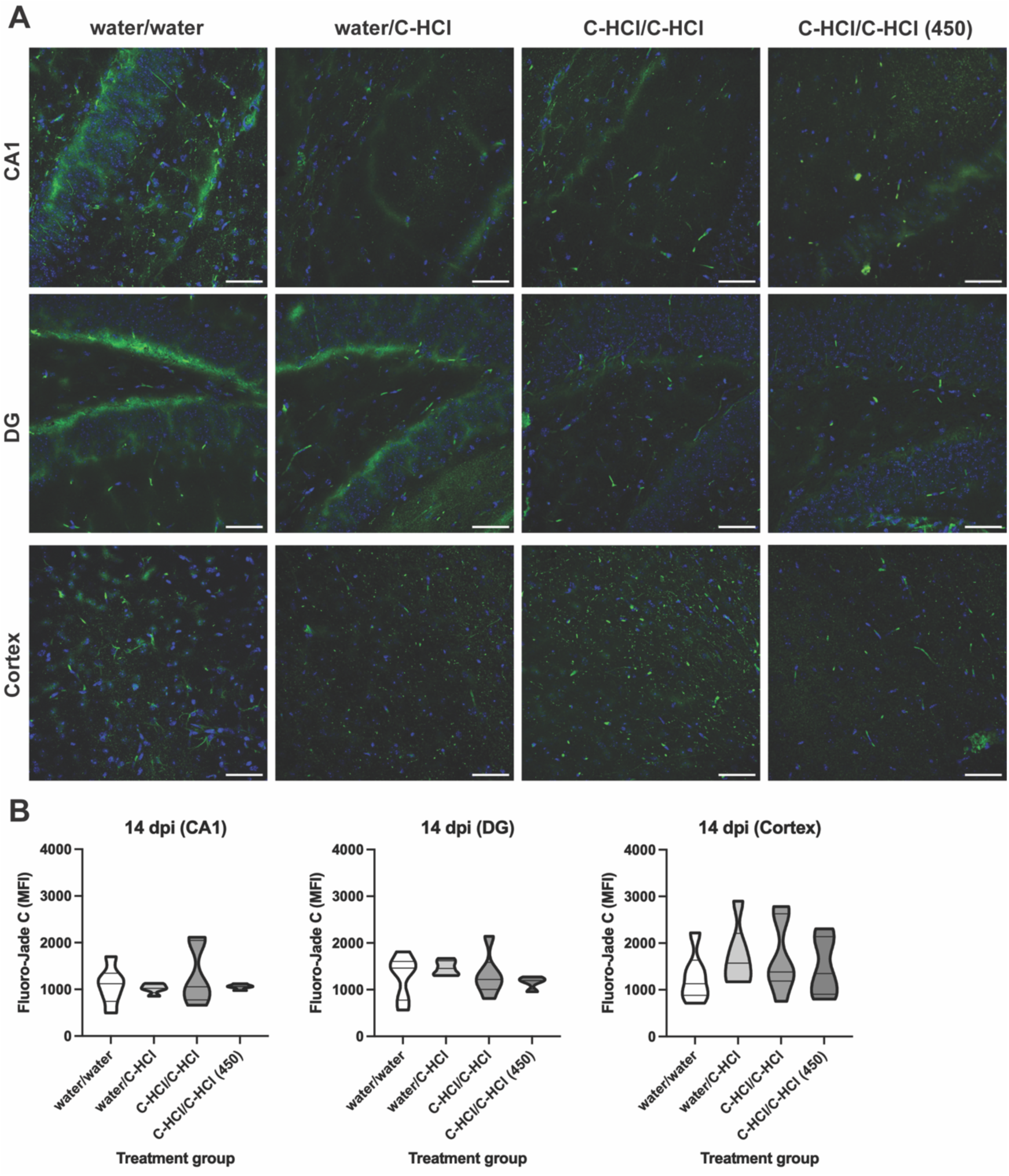
TBI-induced neuron degeneration is unaltered at 14 dpi in female mice treated with C-HCl. (A) Representative immunofluorescent images showing Fluoro-Jade C staining marking degenerating neurons in coronal brain sections from female mice treated with water/water, water/C-HCl, C-HCl/C-HCl, and C-HCl/C-HCl (450) for 7 days before/14 days after TBI. C-HCl concentration was 100 mg/kg unless noted as 450 mg/kg. Objective/scale bar: 40x/50 μm. (B) Quantification of the mean intensity of the fluorescent staining in the CA1 hippocampus (left panel), DG hippocampus (middle panel), and parietal cortex (right panel). Data represented as a volcano plot of *n* = 4-6 per treatment group. No significant differences detected between treatment groups (one-way ANOVA).

### Synapses

Preserving and regenerating synapses is considered a primary goal in neuroprotective therapies. For the analysis of excitatory and inhibitory synapses on mouse brain sections, specific antibodies against synapse-related markers were used as follows: 1) VLUT1 and PSD95 served as markers for excitatory presynaptic and postsynaptic elements, respectively, and 2) VGAT and gephyrin served as markers for inhibitory presynaptic and postsynaptic elements, respectively. No significant changes in either the number of VGLUT+ or PSD95+ puncta were observed across treatment groups in male mice at 14 dpi (**Figure 10**). Similarly, C-HCl treatment did not significantly affect either gephyrin+ puncta numbers in male mice at 14 dpi (**Figure 11**). C-HCl treatment led to a significant increase in the number of VGAT+ puncta in the C-HCl/C-HCl (450) versus C-HCl/C-HCl treated male mice at 14 dpi in the DG region of the hippocampus (**Figure 11**). The male mice in the C-HCl/C-HCl (450) group exhibited significantly reduced VGAT+ puncta numbers compared to the water/C-HCl and C-HCl/C-HCl treatment groups at 14 dpi in the parietal cortex (**Figure 11**). Similar to males, no significant changes in either the number of VGLUT+ or PSD95+ puncta were observed across treatment groups in female mice at 14 dpi (**Figure 12**) apart from significantly lower VGLUT1+ puncta in the hippocampal CA1 region of C-HCl/C-HCl versus C-HCl/C-HCl (450) treated female mice (**Figure 12**). In contrast to males, no significant changes in the number of VGAT+ puncta were observed across treatment groups in female mice at 14 dpi (**Figure 13**). The female mice in the C-HCl/C-HCl (450) group exhibited significantly increased gephyrin+ puncta numbers compared to the water/C-HCl treatment group at 14 dpi in the hippocampal CA1 and compared to the water/water and C-HCl/C-HCl treatment groups at 14 dpi in the hippocampal DG (**Figure 13**).

**Figure 10.**
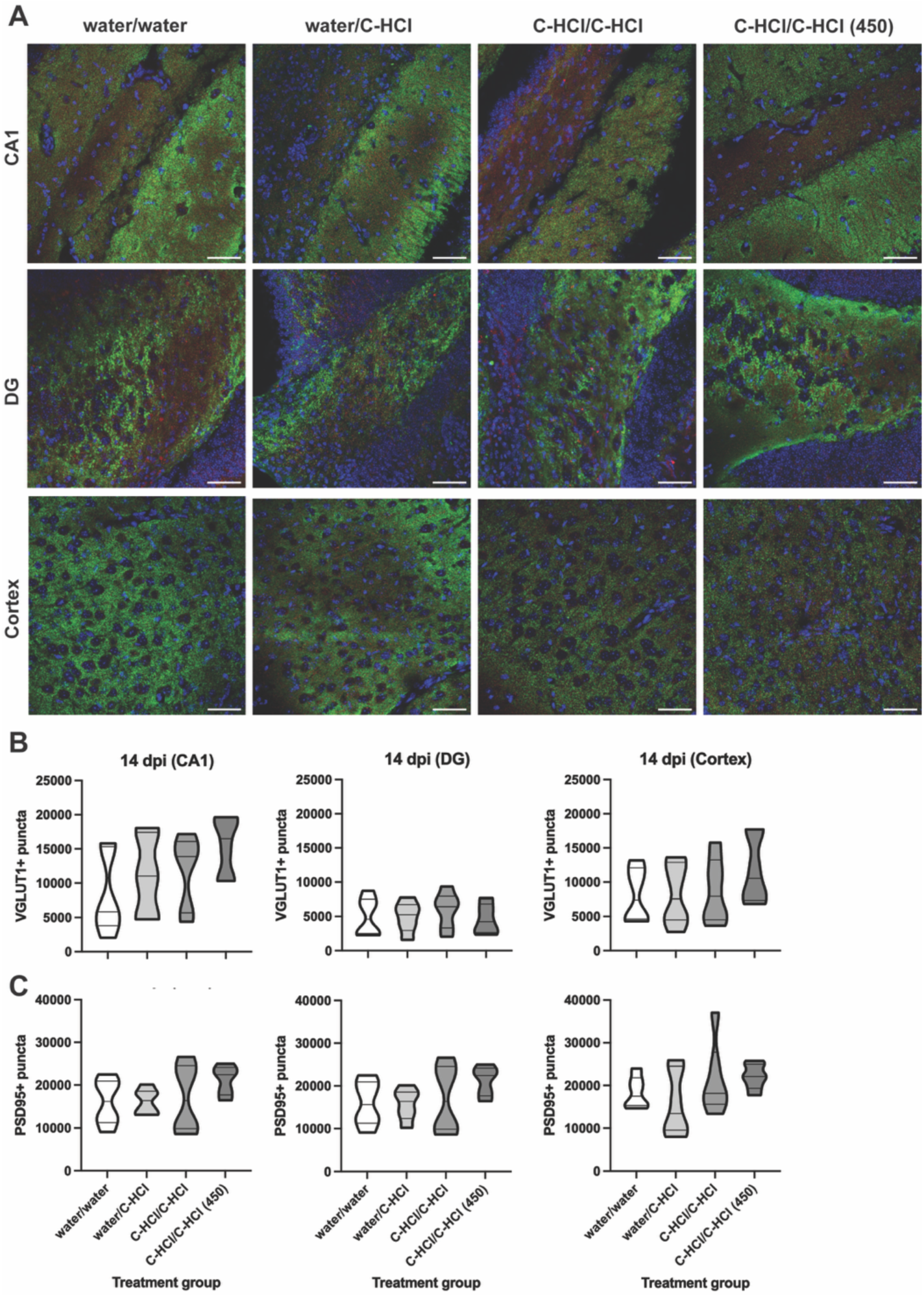
Expression of VGLUT1-positive and PSD95-positive synaptic puncta is unaltered at 14 dpi in male mice treated with C-HCl. (A) Representative immunofluorescent images showing excitatory synapses with VGLUT1 as a marker for excitatory presynapses (green) and PSD95 as a marker for excitatory postsynapses (red) in coronal brain sections from male mice treated with water/water, water/C-HCl, C-HCl/C-HCl, and C-HCl/C-HCl (450) for 7 days before/14 days after TBI. C-HCl concentration was 100 mg/kg unless noted as 450 mg/kg. Objective/scale bar: 40x/50 μm. The total number or (B) VGLUT1-positive synaptic puncta and (C) PSD95-positive puncta were quantified in the CA1 hippocampus (left panel), DG hippocampus (middle panel), and parietal cortex (right panel). Data represented as a volcano plot of *n* = 4-6 per treatment group. No significant differences detected between treatment groups (one-way ANOVA).

**Figure 11.**
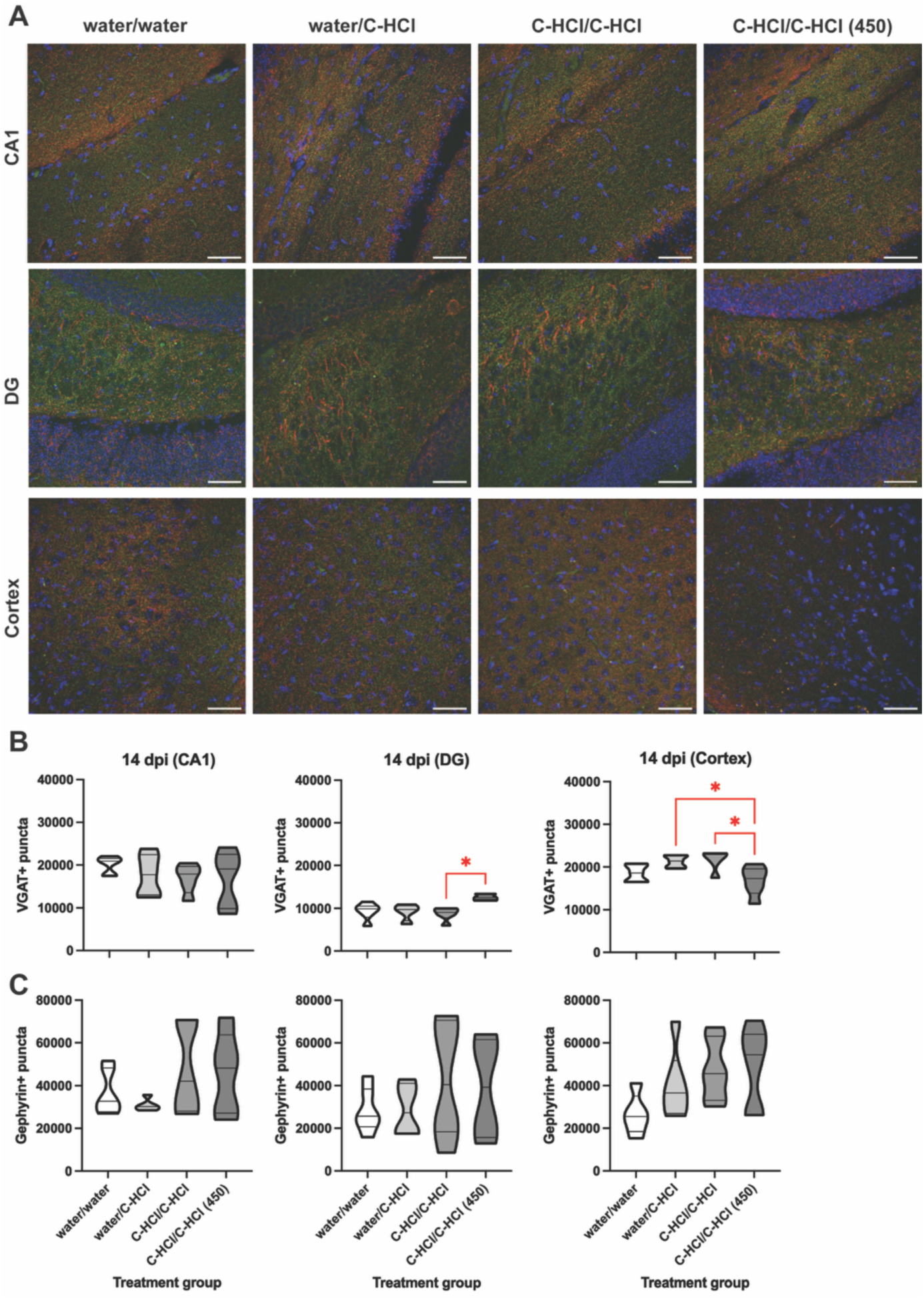
C-HCl alters the number of VGAT-positive synaptic puncta at 14 dpi in male mice in a region-specific manner. (A) Representative immunofluorescent images showing inhibitory synapses with VGAT as a marker for inhibitory presynapses (red) and gephyrin as a marker for inhibitory postsynapses (green) in coronal brain sections from male mice treated with water/water, water/C-HCl, C-HCl/C-HCl, and C-HCl/C-HCl (450) for 7 days before/14 days after TBI. C-HCl concentration was 100 mg/kg unless noted as 450 mg/kg. Objective/scale bar: 40x/50 μm. The total number or (B) VGAT-positive synaptic puncta and (C) gephyrin-positive puncta were quantified in the CA1 hippocampus (left panel), DG hippocampus (middle panel), and parietal cortex (right panel). Data represented as a volcano plot of *n* = 4-6 per treatment group. Statistical significance was determined using a one-way ANOVA followed by Tukey’s multiple comparisons test. \**p* < 0.05.

**Figure 12.**
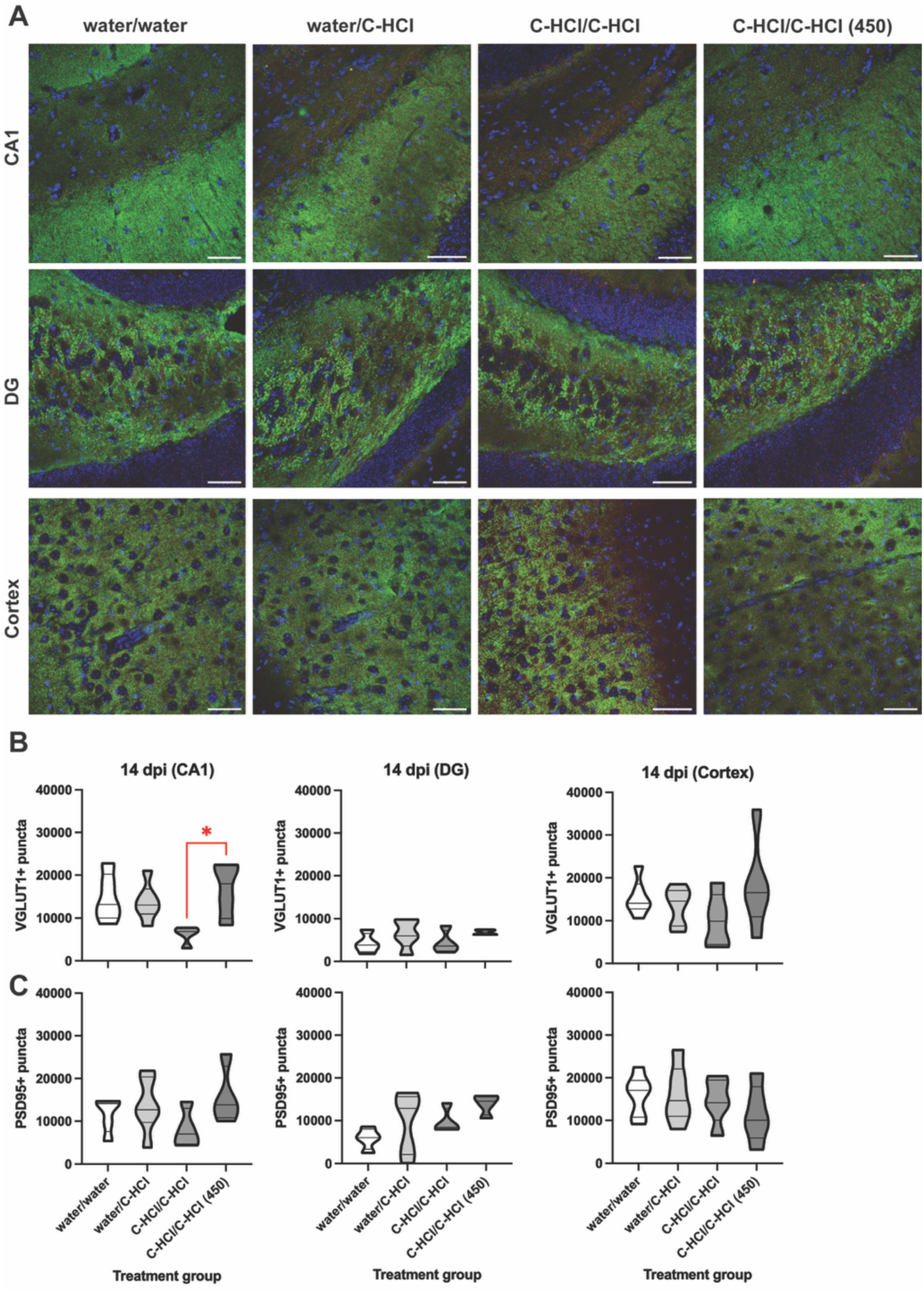
Expression of VGLUT1-positive and PSD95-positive synaptic puncta is unaltered at 14 dpi in female mice treated with C-HCl. (A) Representative immunofluorescent images showing excitatory synapses with VGLUT1 as a marker for excitatory presynapses (green) and PSD95 as a marker for excitatory postsynapses (red) in coronal brain sections from female mice treated with water/water, water/C-HCl, C-HCl/C-HCl, and C-HCl/C-HCl (450) for 7 days before/14 days after TBI. C-HCl concentration was 100 mg/kg unless noted as 450 mg/kg. Objective/scale bar: 40x/50 μm. The total number or (B) VGLUT1-positive synaptic puncta and (C) PSD95-positive puncta were quantified in the CA1 hippocampus (left panel), DG hippocampus (middle panel), and parietal cortex (right panel). Data represented as a volcano plot of *n* = 4-6 per treatment group.

**Figure 13.**
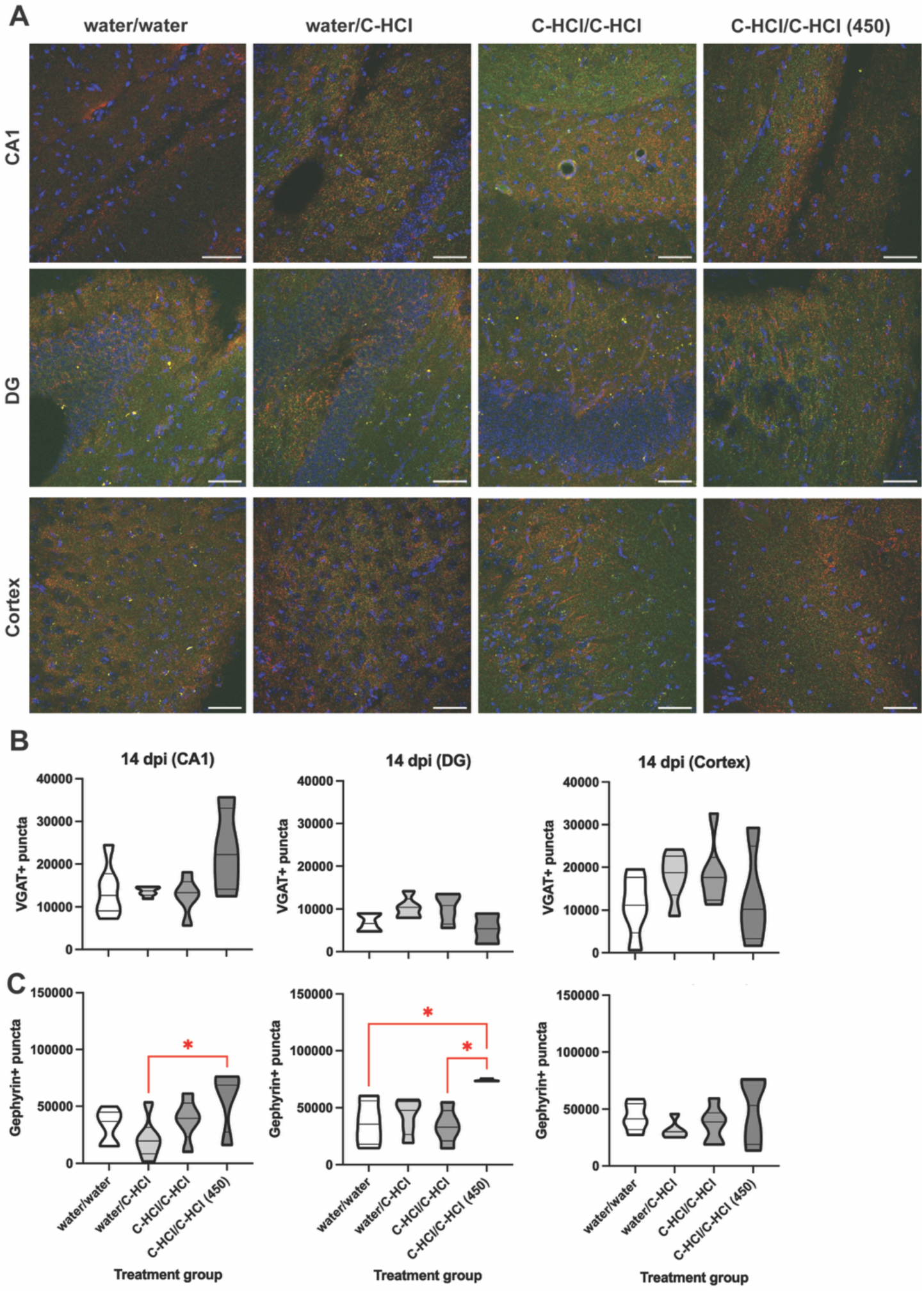
C-HCl increased the number of gephyrin-positive synaptic puncta at 14 dpi in female mice. (A) Representative immunofluorescent images showing inhibitory synapses with VGAT as a marker for inhibitory presynapses (red) and gephyrin as a marker for inhibitory postsynapses (green) in coronal brain sections from female mice treated with water/water, water/C-HCl, C-HCl/C-HCl, and C-HCl/C-HCl (450) for 7 days before/14 days after TBI. C-HCl concentration was 100 mg/kg unless noted as 450 mg/kg. Objective/scale bar: 40x/50 μm. The total number or (B) VGAT-positive synaptic puncta and (C) gephyrin-positive puncta were quantified in the CA1 hippocampus (left panel), DG hippocampus (middle panel), and parietal cortex (right panel). Data represented as a volcano plot of *n* = 4-6 per treatment group.

## Discussion

### Macro-structural injury dynamics and systemic biomarkers

At 14 dpi, exogenous C-HCl treatment did not yield a macro-structural repair, showing no significant impact on gross lesion size, hemispheric tissue loss, or third ventricle expansion in either male or female mice. Concurrently, systemic fluid biomarkers of acute neuroaxonal degradation (NfL, t-Tau) and astrocytic damage (GFAP) remained statistically unaffected by the treatment. However, baseline TBI-induced lesion areas were significantly larger in female mice compared to male mice across all groups. This prominent sex-based divergence in baseline damage reflects established literature characterizing differential vulnerabilities to biomechanical forces and early pathophysiological cascades between male and female rodents ^7,8^. The absence of treatment-induced gross anatomical shifts or changes in late-stage peripheral protein shedding suggests that the neuroprotective action of C-HCl operates downstream of macroscopic cavitation. This aligns with historical models demonstrating that neurometabolic interventions frequently rescue local cellular architecture and bioenergetic viability without fundamentally altering the primary, macro-scale mechanical lesion boundary ^9^.

### Glial activation and neuroinflammation

Immunohistochemical analysis revealed a distinct, sex-specific anti-inflammatory effect localized predominantly within the hippocampal CA1 region of female mice. The higher dose C/C-HCl regimen significantly blunted microglial activation (Iba1 MFI) relative to water controls and reduced both astrocyte density and activation (GFAP % area and MFI) in female mice. Conversely, reactive gliosis in male mice remained unaltered by treatment at 14 dpi. These findings directly complement the growing understanding of creatine as a metabolic buffer capable of modulating energy-demanding inflammatory cascades. In the context of TBI, reactive astrocytes and microglia experience severe bioenergetic strain to maintain ion homeostasis and drive cytokine production. By stabilizing intracellular ATP pools, creatine appears to dampen this hyper-reactive state ^9^. Crucially, the restriction of this anti-inflammatory effect to female CA1 regions highlights a metabolic or hormonal gatekeeping mechanism. This suggests that the capacity of C-HCl to suppress chronic gliosis is highly dependent on sex-stratified cellular environments or baseline neurochemical profiles ^10^.

### Neuronal survival and neuroprotection

In direct contrast to the glial results, microstructural neuroprotective effects of C-HCl were heavily skewed toward male mice. C/C-HCl (450) and W/C-HCl regimens significantly reduced the number of degenerating Fluoro-Jade C+ neurons within the male hippocampal CA1 and DG regions. Female mice showed no significant treatment shifts in neurodegeneration. This localized reduction in Fluoro-Jade C staining supports wide-ranging literature characterizing creatine as a classic neuroprotective agent capable of mitigating secondary cell death. The preservation of hippocampal neurons in males, but not females, underscores a complex inversion of target mechanisms between sexes. While females benefited via upstream glial stabilization, males derived downstream advantages through direct neuronal survival pathways, reinforcing the need for sex-stratified analysis in neurometabolic therapies. The localized preservation of hippocampal neurons in males validates long-standing preclinical literature identifying creatine as a classic neuroprotective countermeasure against secondary metabolic failure ^9^. TBI routinely triggers an excitotoxic energy crisis characterized by mitochondrial depolarization and intracellular calcium overloads. Creatine supplementation mitigates this decay by maintaining mitochondrial membrane integrity and optimizing bioenergetics^9^. The paradoxical inversion of targets between the sexes, where females demonstrated upstream glial stabilization but males exhibited direct downstream neuronal survival underscores a complex, sex-dependent therapeutic mechanism that requires explicit stratification in translational TBI models.

### Inhibitory Synaptic Remodeling & Circuit Plasticity

While sweeping changes in excitatory synaptic markers (VGLUT1 and PSD95 puncta) were largely absent, C-HCl treatment drove profound, targeted alterations in inhibitory synaptic machinery (VGAT and gephyrin). However, the molecular strategies deployed were distinct. Males demonstrated presynaptic modulation, marked by a localized increase in presynaptic GABAergic transport (VGAT+ puncta) within the hippocampal DG, alongside a simultaneous decrease in the parietal cortex under the high-dose C-HCl (450) regimen. Conversely, females exhibited structural postsynaptic modulation, driving significant increases in scaffolding elements (gephyrin+ puncta) in both the CA1 and DG regions. Post-traumatic circuitry is heavily defined by a breakdown of recurrent network inhibition, a pathological hallmark that directly precipitates post-traumatic epilepsy (PTE) and cognitive deterioration ^11^. These findings indicate that creatine aids in restoring network balance by selectively fortifying the GABAergic framework. Preclinical literature confirms that delayed or chronic creatine administration counteracts TBI-induced losses of GABAergic functional components ^12^. The novelty of our current data lies in its mechanistic divergence: males primarily adjust the presynaptic vesicular supply lines (VGAT), whereas females reinforce the postsynaptic receptor-anchoring architecture (gephyrin) ^13^. This implies that while the ultimate therapeutic goal (enhanced circuit inhibition) is uniform, the underlying molecular pathways affected by C-HCl treatment in the context of TBI are dictated by biological sex.

### C-HCl

The effects of creatine on cellular energy reserve, mitochondrial function and inflammation ^1,14,15^ supports its potential use in TBI. Indeed, studies examining creatine supplementation on TBI in pre-clinical animal models have reported improvements ranging from significant reductions in cortical damage to reductions in brain lactate levels following creatine administered in food or given as ip injection prior to injury ^9,16^. In these studies, creatine monohydrate, was used prophylactically at relatively high doses (1% of the daily diet ^16^ or 3000 mg/kg ip injection ^9^). An open label trial by Sakellaris and colleagues ^17^ administered a daily dose of creatine monohydrate (400 mg/kg or approximately 20 g per day) to adolescents with TBI and reported improvement in cognitive and behavioral measures. Such findings are in line with the 20-30 g daily doses of creatine monohydrate being used for neurodegenerative diseases ^15,18^. While these studies suggest high dose creatine supplementation may provide prophylactic and treatment advantages for recovery from TBI, the high-volume loading phase and large doses required limit extended use and effectiveness.

Creatine HCl, is a newer form of creatine with increased aqueous solubility and bioavailability compared to creatine monohydrate ^19^. Both clinical case reports and clinical trials suggest creatine HCl dosing, especially for applications involving brain related outcomes, is significantly reduced compared to standard creatine monohydrate ^20^. Given the effectiveness of creatine HCl, the present study examined creatine HCl supplementation in various pretreatment and post-treatment regimens at doses that are more clinically realistic. To put the current dosage regiments in context, using the FDA-based guidelines for determining human equivalent dose, the doses used in the present study ranging from 100 mg/kg to 450 mg/kg would represent approximately 0.5 g and 2 g daily human equivalent doses. The FDA guidelines are intentionally conservative estimates, thus using allometric dosing calculations accounting for both surface area and metabolic differences puts the estimated human dose equivalents at 1.5 g and 6 g ^21^. While confirmational studies are required, the doses of creatine hydrochloride supplementation used in the current study represent a substantial reduction from previous studies using creatine monohydrate. The findings of the present study suggest C-HCl supplementation holds promise as a neuroprotective therapeutic strategy for TBI.

## Conclusion

In summary, this study demonstrates that while a 14-dpi regimen of exogenous C-HCl does not alter macro-structural lesion boundaries or systemic fluid biomarker profiles (NfL, t-Tau, GFAP), it exerts a potent, highly localized neuroprotective influence within the hippocampus. Crucially, this therapeutic window is governed by a profound biologically sex-dependent split in target pathways. Rather than operating uniformly across both sexes, the mechanism of action of C-HCl diverges into distinct cellular domains. Females derive primary benefit via an upstream glial stabilization cascade, marked by a robust reduction in reactive astrogliosis and microglial activation within the hippocampal CA1 region, accompanied by the structural reinforcement of postsynaptic inhibitory elements (gephyrin). Males bypass glial modification entirely, instead showing a direct downstream preservation of cellular survival pathways (reduced Fluoro-Jade C+ neuronal degeneration in the CA1/DG) alongside an optimization of presynaptic inhibitory machinery (VGAT). Ultimately, these data indicate that C-HCl successfully counteracts chronic secondary injury cascades, mitigating neurodegeneration, dampening neuroinflammation, and remodeling inhibitory circuitry to prevent post-traumatic network hyperexcitability. However, the molecular strategies deployed to achieve this functional homeostasis are entirely distinct between male and female mice. These findings underscore a critical translational insight: neurometabolic interventions for TBI are not one-size-fits-all, and preclinical therapeutic pipelines must explicitly stratify mechanistic targets by biological sex to maximize efficacy. Future studies aimed at determining the optimal dose of C-HCl for neuroprotection following TBI in male and female mice will be essential. To put the current dosage regiments in context, there are two different human equivalent approaches; (i) one uses the FDA-based guidelines for determining human equivalent dosing, and the doses used in the present study ranging from 100 mg/kg to 450 mg/kg would represent approximately 0.5 g and 2 g daily human equivalent doses. However, FDA guidelines are intentionally conservative estimates, thus (ii) using allometric dosing calculations accounting for both surface area and metabolic differences puts the second estimated human dose equivalents at 1.5 g and 6 g ^21^. While confirmational studies are required, the doses of creatine hydrochloride supplementation used in the current study represent a substantial reduction from previous studies using creatine monohydrate. The findings of the present study suggest C-HCl supplementation holds promise as a neuroprotective therapeutic strategy for TBI. And while behavioral outcomes were not formally assessed in the present study, mice receiving creatine supplementation in general were more active post injury suggesting a more quantitative assessment of behaviors is also warranted.

## Acknowledgements

We recognize the use of the University of Nebraska Medical Center – Tissue Science Facilities, RRID:SCR_012465, which receives support from the Buffett Cancer Center Support Grant P30 CA036727 and the UNMC Department of Pathology, Microbiology, and Immunology.

## Author Disclosure Statement

The authors have declared no competing interest.

