## Supplemental Figure S1 for "Sex-Dependent Effects of Creatine Hydrochloride on Neuroinflammation, Neuronal Survival, and Inhibitory Synaptic Remodeling Following Traumatic Brain Injury in Mice"

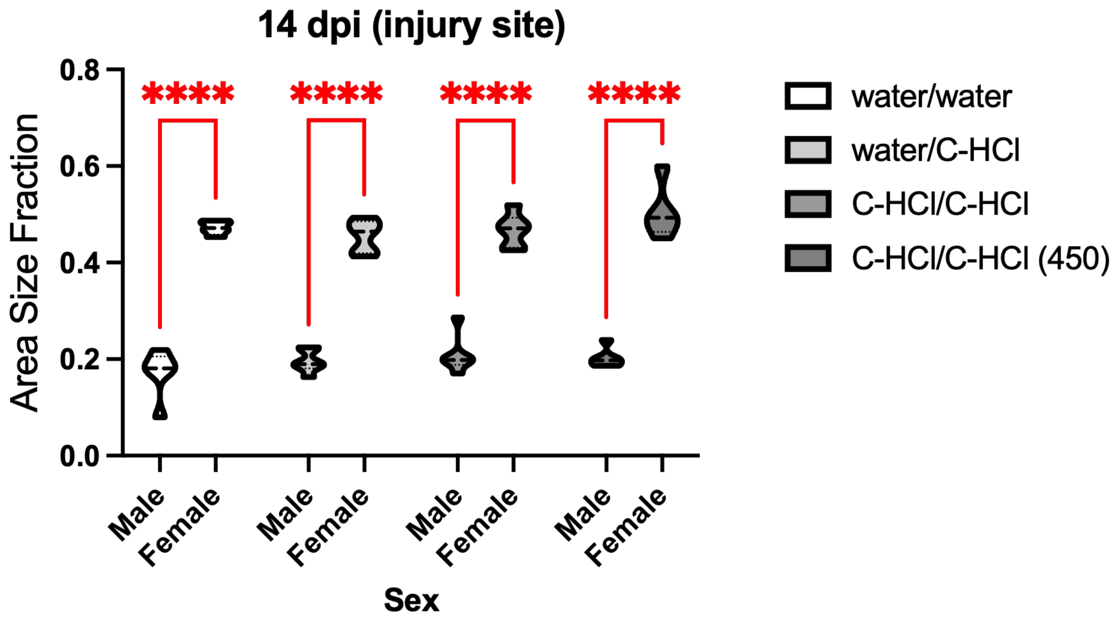


**Supplemental Figure S1. TBI-induced tissue damage is greater in female mice.** The area fraction of the injury site (area of the injury site/area of the contralateral brain hemisphere) for male and female mice at 14 dpi. Data represented as a volcano plot of *n* = 5-6 per treatment group. Statistical significance was determined using a two-way ANOVA followed by Sidak's multiple comparisons test. ****$p$ < 0.0001.
